# Dynamic causal modelling of fluctuating connectivity in resting-state EEG

**DOI:** 10.1101/303933

**Authors:** Frederik Van de Steen, Hannes Almgren, Adeel Razi, Karl Friston, Daniele Marinazzo

## Abstract

Functional and effective connectivity are known to change systematically over time. These changes might be explained by several factors, including intrinsic fluctuations in activity-dependent neuronal coupling and contextual factors, like experimental condition and time. Furthermore, contextual effects may be subject-specific or conserved over subjects. To characterize fluctuations in effective connectivity, we used dynamic causal modelling (DCM) of cross spectral responses over 1 min of electroencephalogram (EEG) recordings during rest, divided into 1-sec windows. We focused on two intrinsic networks: the default mode and the saliency network. DCM was applied to estimate connectivity in each time-window for both networks. Fluctuations in DCM connectivity parameters were assessed using hierarchical parametric empirical Bayes (PEB). Within-subject, between-window effects were modelled with a second-level linear model with temporal basis functions as regressors. This procedure was conducted for every subject separately. Bayesian model reduction was then used to assess which (combination of) temporal basis functions best explain dynamic connectivity over windows. A third (between-subject) level model was used to infer which dynamic connectivity parameters are conserved over subjects. Our results indicate that connectivity fluctuations in the saliency network comprised both subject-specific components and a common component. For the default mode network, connectivity trajectories only showed a common component. For both networks, connections to higher order regions appear to monotonically increase during the one minute period. These results not only establish the predictive validity of dynamic connectivity estimates – in virtue of detecting systematic changes over subjects – they also suggest a network-specific dissociation in the relative contribution of fluctuations in connectivity that depend upon experimental context. We envisage these procedures could be useful for characterizing brain state transitions that may be explained by their cognitive or neuropathological underpinnings.

## Introduction

Evolution over time of segregated and integrated brain activity is somehow intrinsic to its definition in terms of distributed neuronal dynamics. On the other hand the genesis, nature, and the time scale of these changes are diverse, and represent an increasing focus of research. Structurally, the rewiring of white matter in humans – through axonal growth or pruning – is unlikely to occur during late development (i.e. between age 2 and 18); however, changes in white matter tracts, such as increased (possibly activity dependent) myelination and axonal diameter have been shown by several studies (e.g. Giedd et al., 1999; Paus, 2010; Rademacher et al., 1999). Together with these structural changes, long and short range functional connectivity (FC) also changes during development (Hagmann et al., 2010). Here, FC is defined as the statistical dependencies among observed neurophysiological measures (e.g. correlation between blood oxygen level dependent (BOLD) signals; Friston, 2011). More recently, changes in FC at a shorter time-scale have been investigated (e.g., Allen et al., 2014; Chang and Glover, 2010; Vidaurre et al., 2017; Wang et al., 2016). The time scale of these studies is on the order of minutes (i.e. during a typical resting-state functional magnetic resonance imaging (rs-fMRI) protocol) – as opposed to the developmental studies which are in the order of years. Typically, the dynamic FC is quantified using a (sliding) window approach: the resting state time-series are segmented into (partially overlapping) windows and FC is calculated for each window. This approach allows researchers to assess the trajectory of FC over time for different networks/states. For example, Vidaurre et al. (2017) showed, using a hidden Markov model, that the transitions between networks (states) are non-random. Two recent studies (Heitmann and Breakspear, 2017; Liégeois et al., 2017) also made the important point of distinguishing the meaning of “dynamic” from the statistical point of view (i.e., non-uniformity in time) from the link of these fluctuations to the intrinsic dynamics of neural populations that generate the recordings under examination.

However, studies dealing with fluctuations in functional connectivity cannot directly provide evidence of fluctuations in the underlying effective connectivity – which is defined as the directed (causal) influence one neuronal population exerts on the other. Effective connectivity can only be inferred using a forward or *generative model* of how activity in one brain region, or external stimulus, affects activity (i.e. causes a change in activity) in another brain region. Since we usually do not measure neural activity directly, the generative model usually includes a forward model of how (hidden) neural activity is mapped to the (observed) measurement (Friston, 2011). Inferring effective connectivity therefore requires the inversion of this generative model. This can be done with standard (variational) Bayesian model inversion, as in dynamic causal modelling (DCM; David et al., 2006; Friston et al., 2012, 2003).

Several studies have investigated fluctuations of effective connectivity obtained from electrophysiological recordings (Cooray et al., 2015; Papadopoulou et al., 2017, 2015) and fMRI (Park et al., 2017) using DCM. These electrophysiological studies were conducted in the context of tracking connectivity changes around periods of epileptic events, while the fMRI study investigated healthy subjects during rest. Cooray et al. (2015) used Bayesian belief updating, in which between window differences were modelled as a random walk. This contrasts with the approach in Papadopoulou et al. (2015), where the authors used a set of temporal basis functions to model connectivity fluctuations. In this approach, all windows were inverted simultaneously, making it computationally expensive. In the current work, a hierarchical model is employed: each window is inverted independently and subsequently fluctuations are modelled over windows; i.e., at a between-window (within-subject) level. More specifically, a Bayesian linear model with temporal basis functions is used to model between window-differences. Parametric empirical Bayes (PEB, Friston et al., 2016) is the procedure used to estimate this second level model. The main difference between Papadopoulou et al. (2017) and Park et al. (2017) is that the latter used multilevel PEB to make inferences at the group level, while the former concatenated the data from different subjects at the group level. In this paper, we use the same technology to assess fluctuations in effective connectivity at the within and between-subject level, with a special focus on systematic, time sensitive fluctuations that may be conserved over subjects.

In this work, we used eyes-open resting state electroencephalographic data (EEG) time series from healthy subjects. Our goal was to quantify how effective connectivity fluctuates over time using hierarchical Bayesian modelling. More specifically, we investigate whether connectivity fluctuates in a subject-specific manner and/or whether there are systematic components embedded within the connectivity trajectories that are conserved over subjects. We addressed this by using DCM for cross spectral density data features (CSD) combined with (multilevel) PEB (Friston et al., 2016, 2012). DCM was used to infer effective connectivity from windowed resting-state EEG time-series. Then, PEB was employed to characterize the trajectory of DCM parameters during the 1 min recording session. Applying the approach described in Park et al. (2017) to EEG allowed us to track fluctuations in DCM connectivity parameters on the time-scale of seconds and minutes. It is important to note that the generative model (i.e., DCM) for electromagnetic time series is biophysically more detailed compared to DCM for rs-fMRI: the hidden state in DCM for CSD applied to EEG consist of voltages and currents of specific neuronal cell populations, while in fMRI the hidden state are an abstraction of neural activity (Friston et al., 2014; but also see recent developments Friston et al., 2017).

Our hypothesis was that we would be able to detect within subject (between window) fluctuations – in all subjects – in key (intrinsic) brain networks and, crucially, some of these fluctuations would have predictive validity in the sense that they would show systematic time effects in relation to the onset of the recording session. To address this hypothesis, we therefore analyzed a large (publicly available) cohort of data, paying special attention to the hierarchical (PEB) modelling of random effects at the within-window, within-subject and within-group levels respectively. This analysis is offered as a proof of principle that systematic aspects of dynamic effective connectivity can be recovered from EEG data. We anticipate that the methods described below may be usefully applied to test for experimental effects on dynamic connectivity and implicit short-term synaptic plasticity.

## Methods

### Data and pre-processing

In this study, we used one minute eyes open EEG recordings from the EEG Motor Movement/Imagery PhysioNet dataset (Goldberger et al., 2000; Schalk et al., 2004). The data was acquired using the BCI2000 system (www.bci2000.org). The EEG channels were placed on the scalp according to the international 10-10 system (Chatrian et al., 1985). The data was provided in EDF+ format, containing 64 EEG channels, each sampled at 160 Hz.

The data were preprocessed using EEGLAB running on MATLAB (Delorme and Makeig, 2004). The 60Hz power line noise was first removed using the Cleanline EEGLAB plugin. Afterwards, the data was high-pass filtered using default settings, with a lower-cutoff of 1Hz. Then, a low-pass filter with high-cutoff of 45 Hz and default settings was applied. Periods of data contaminated with blink artifacts were repaired using independent components analysis. Bad channels were removed based on visual inspection. Finally, the data was average-referenced.

### Dynamic causal modelling

The preprocessed data was imported in SPM12 (Wellcome Trust Centre for Human Neuroimaging; www.fil.ion.ucl.ac.uk/spm/software/spm12). DCM for CSD was used for further analyses (DCM12, Friston et al., 2012). In brief, DCM explains the observed CSD by combining a generative (biophysically plausible) neural mass model and an observation model that maps neuronal states to the observed data. Each electromagnetic source or node is equipped with three neuronal subpopulations: pyramidal cells, inhibitory interneurons and spiny stellate cells (the ‘ERP’ model; Moran et al., 2013). The connections between subpopulations within a node are termed intrinsic connections, while connections between nodes are termed extrinsic connections. There are three types of extrinsic connections: forward, backward and lateral connections, which differ in terms of their origin and target subpopulation. The type of connection can be determined based on the hierarchical organisation of the cortex (Felleman and Van Essen, 1991). The default forward model provided by SPM12 was used for the observation equation (which is in essence a linear mapping from the hidden neural states to the EEG sensor spectral densities; i.e., a conventional lead field or gain matrix).

In DCM for EEG, the anatomical locations of the nodes need to be specified *a priori.* We inverted, two (fully) connected models (hierarchical inversion, see below), one for the default mode network and one for the saliency network. For the default network, the following four nodes were chosen: left and right lateral parietal area (l/rLP; MNI coordinates: −46 −66 30; 49 −63 33, respectively), posterior cingulate/Precuneus (Prec; MNI coordinates: 0 −58 0 and medial prefrontal cortex (mPFC; MNI coordinates: −1 54 27, see Fig 1.A). For the saliency network, left and right lateral parietal (l/rLP; MNI coordinates: ±62 −45 30) area, left and right anterior prefrontal cortex (l/raPFC; MNI coordinates: −35 45 30; 32 45 30) and dorsal anterior cingulate cortex (dACC; MNI coordinates: 0 21 36 see Fig 1.A) were specified. These nodes were chosen based on the sources used in (Razi et al., 2017). In both networks, the nodes were connected with forward, backward and lateral connection as described by (David et al., 2006, 2005; Felleman and Van Essen, 1991). See Fig 1.B for a schematic presentation of the presumed coupling among the sources. Each node was treated as a patch on the cortical surface (‘IMG’ option in SPM12).

**Figure 1.**
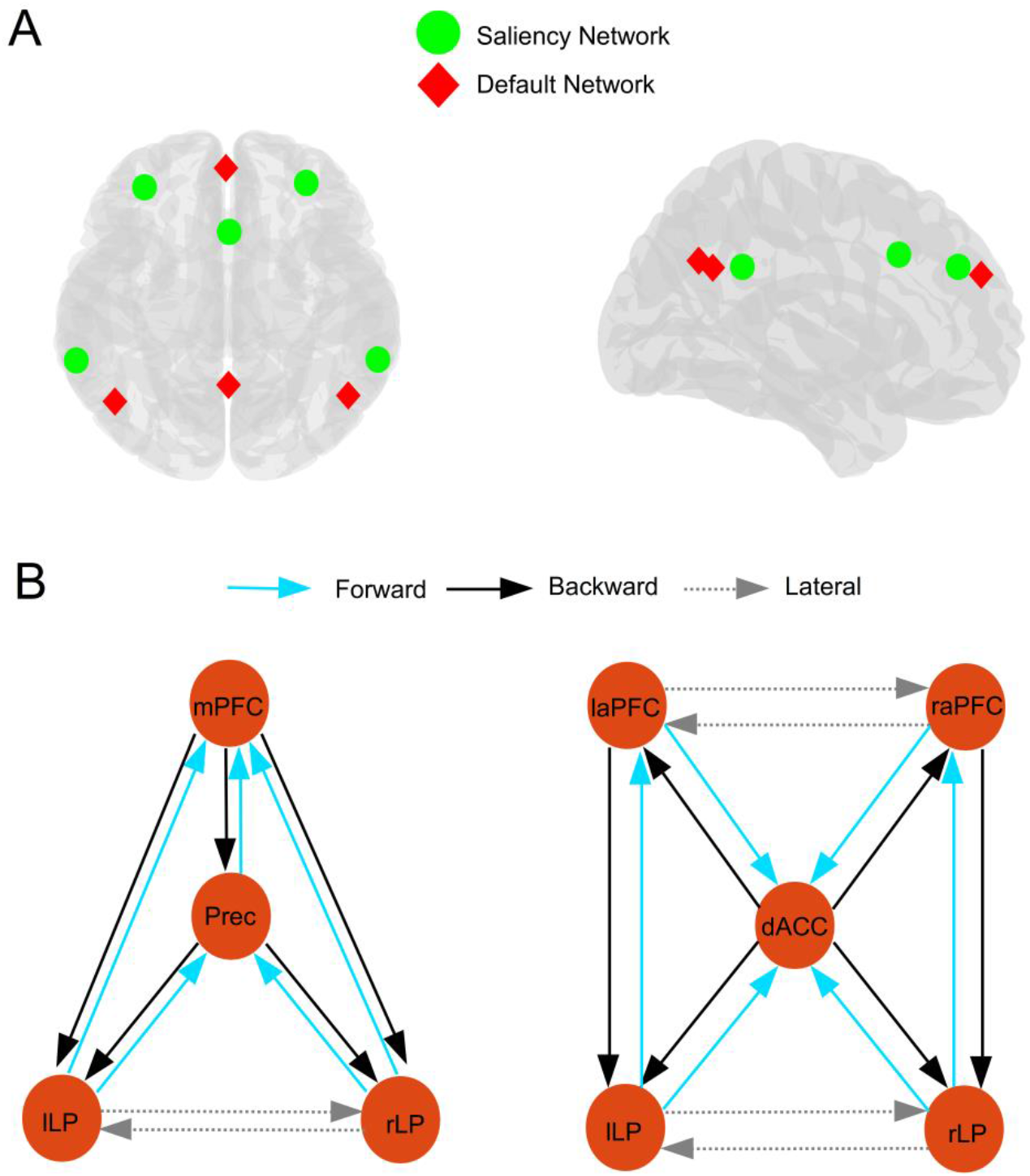
The default and saliency network studied in this paper are shown. Panel A shows the locations of sources included in both network. The default mode network included bilateral parietal areas (l/rLP), medial prefrontal cortex (mPFC) and the precuneus (Prec). The saliency network included bilateral parietal areas (l/rLP), dorsal anterior cingulate cortex (dACC) and bilateral anteriorprefrontal cortex (aPFC). Panel B shows a schematic representation of the fully connected model in both networks. Light blue arrows depict forward connections, black arrows depict backward connections, and gray arrows depict lateral connections. The left panel corresponds to the default network while the right panel represents the saliency network.

### Time-varying DCM using Parametric Empirical Bayes

To characterize dynamic effective connectivity, the resting state EEG time-series (one minute duration) were divided into 1 second consecutive epochs (Papadopoulou et al., 2017, 2015). Foreach time-window, two DCM’s (one for each network) were fitted to the data as described above. Then, the between-window differences within a network were modelled using a second level (general) linear model with a set of temporal basis functions (i.e., a discrete cosine set and a monoexponential decay function) as regressors. More specifically, a hierarchical generative model was employed in which first level (i.e. window-level) DCM parameters are treated as a linear mapping from a second level model (i.e. subjects-level):

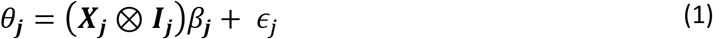

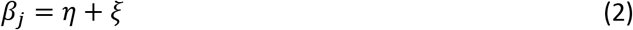

Here, 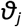 are the vectorised (multivariate) DCM parameters. More specifically, the first Z = 1,…,B elements of 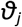 are the DCM parameters of the first window. The vectorised DCM parameters are thus stacked window-wise with B equals the number of DCM parameters and I = 1,…,W where W equals the number of windows so that *θ_j_* = [*θ*_11*j*_,…, *θ*_1*Bj*_,…,*θ*_*W*1*j*_,…,*θ_WBj_*,]^*T*^. ***X**_j_* refers to the second level design matrix of the jth subject. The first column is a constant term while the other columns are temporal basis functions.

In this work we specified five temporal basis functions. These include a discrete cosine transformation (DCT) basis set, and one mono-exponential decay basis function (see Fig. 2). This second level design matrix describes how the DCM parameters change over time. The Kronecker tensor product (⊗) of ***X**_j_* and ***I**_j_* (the identity matrix of size B) means that each within window, each parameter can show one or more second level effects. Furthermore, *β_j_* are the second level parameters represented as a column vector of length BW. The last term in (1), *ε_j_*, is the interwindow variability (i.e., random effects) and *ξ* corresponds to the amplitude of random fluctuations in the second level parameters.

**Figure 2.**
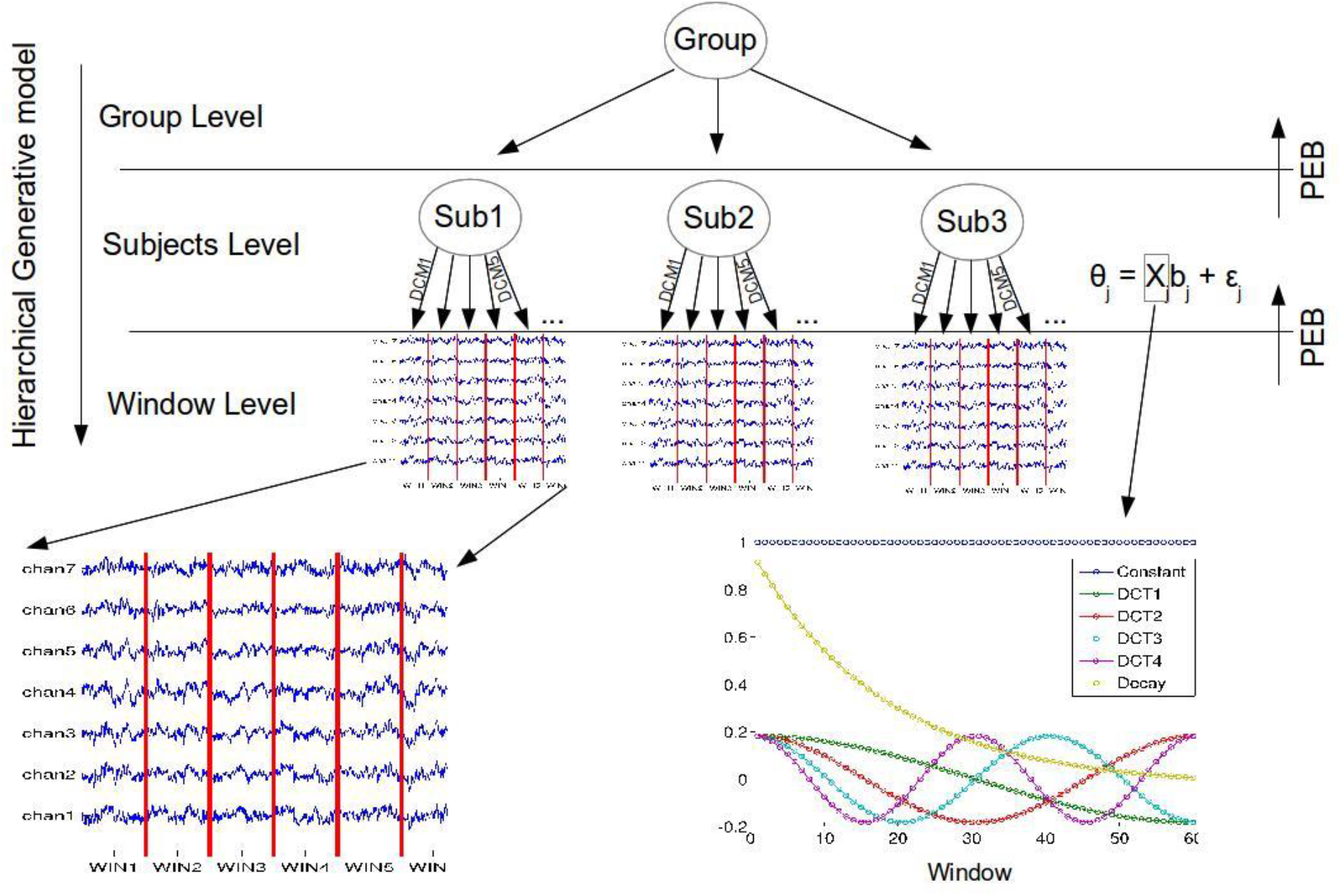
The hierarchical generative model used in this work. The lowest level depicts the window-level. At this level, two DCMs – one for the default mode network and one for the saliency network – were inverted for each window. Using parametric empirical Bayes (PEB), time dependent differences between windows are modelled at the within-subject level. The temporal basis functions used to model these differences are shown in the lower right panel. At the group level, the mean of time-dependent changes in effective connectivity was assessed, using PEB of between-subject effects (group-PEB).

Using PEB framework, the parameters in (1) and (2) can be estimated for each subject (i.e., subject-specific PEB) from the posterior means and covariances from each window as described in (Friston et al., 2016). Next, Bayesian model reduction can be used to efficiently estimate reduced second level models, from the full second level model (see Friston and Penny, 2011; Friston et al., 2016, for more details). Here, we derived all combinations of the five temporal basis functions – so that for each subject, 32 reduced models were estimated from the full second level model (using spm_dcm_bmc_peb.m). Bayesian model comparison of those second level models was conducted by summing the log evidences for each second level model over subjects.

Finally, the second level parameter estimates (i.e., posterior mean and covariance), from the second level model were entered into a third level (PEB) model for group-level inference. Specifically, subject-specific parameters of between-window effects for each DCM parameter were modelled with a second level model but now, the design matrix contained only a constant term. In other words, the average of – or conserved – between-window effects across subjects were modelled (we refer to this as the group-PEB). This enabled us to ask which between-window effects of specific connections are conserved over subjects. Finally, any parameters of the group-PEB that do not contribute to the log evidence were pruned away, using Bayesian model reduction and a greedy search (implemented in spm_dcm_peb_bmc.m).

To preclude local minima at the first (within-window) level, we used PEB scheme for estimating the within-window DCM parameters. In this application of PEB, the first level parameters are iteratively re-estimated under the assumption they are generated from a between-window mean (using spm_dcm_peb_fit.m). This uses empirical shrinkage priors to pull solutions away from local minima (Friston et al., 2015). Since our study involved inverting a large number of DCMs, the spm_dcm_peb_fit.m code was compiled as a stand-alone executable (using the MATLAB mcc command). This enabled the initial estimation to be conducted in parallel on the Ghent University High Performance Computing infrastructure.

In summary, Bayesian model reduction (i.e., selection of reduced models) of second level (subject-specific) models was used to identify the temporal basis functions that best explained between-window differences. The group-PEB analysis was then performed to assess which time-dependent components areconservedoversubjects.

## Results

Our goal was to estimate time-dependent changes in effective connectivity using rest-EEG. More specifically, we wanted to characterize the connectivity trajectories that may or may not be conserved over subjects. Segmented data was analysed with DCM for CSD combined with (multilevel) PEB. We first estimated a full second level PEB model for each subject. Then using Bayesian model reduction, all combinations of second level regressors (except the constant, which was always included in the model) were derived from the full model (see Fig. 3A for the second level model space). The most likely combinations of regressors were detected by taking the sum of the free energies over subjects, for all combinations of second level basis functions. The results of these analyses are shown in Fig. 3B. It is clear that for the saliency network, the inclusion of all basis functions appear to be necessary for explaining the between-window differences in effective connectivity (across subjects). For the default mode network, we observe that only the monotonic decay was relevant to model the between window differences.

**Figure 3.**
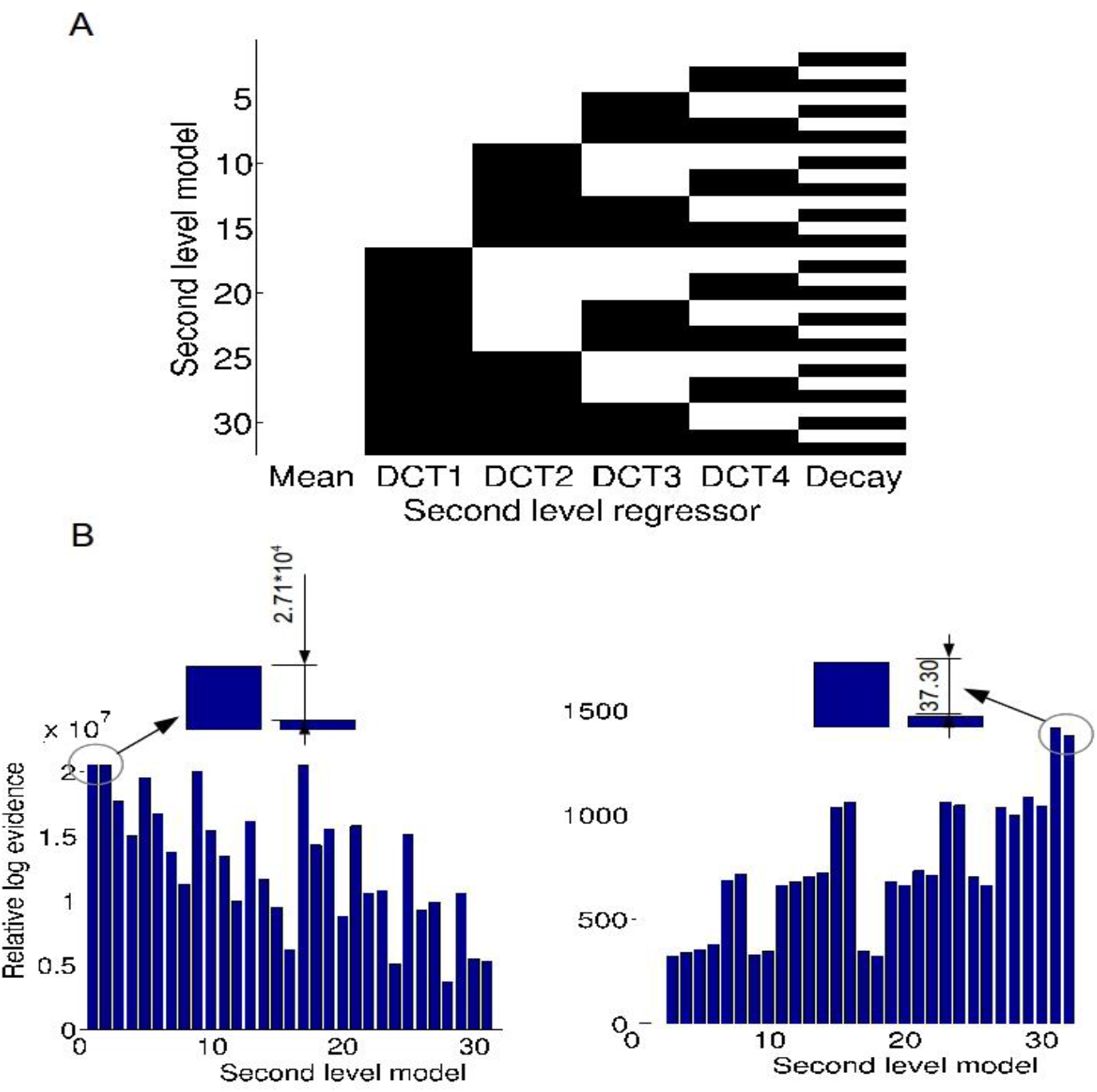
The results for the second level (between-window, within-subject) Bayesian model comparison. Panel A shows the second level model space. These included all combinations of second level basis functions. The constant term was included in all reduced models. Panel B shows the sum of the free energies (over subjects), relative to the least evident model for all reduced second level models for the saliency network (left) and default network (right).

Next, we performed a group PEB of second level effects to investigate which DCM parameter(s) show one or more time dependent effects that are conserved across subjects. This means that for the group PEB, there are 6 (basis functions + constant) by 12 (DCM) for the default network and 6 by 16 parameters for the saliency network. Note that we only modelled extrinsic connections. These are parameterized in terms of log gains. This way, the connections are constrained to be excitatory (extrinsic or between source cortico-cortical connections in the brain are excitatory or glutamatergic). In the neural model, intrinsic connectivity is modelled with four between subpopulations couplings (for each source separately), where contextual changes in intrinsic coupling are modelled by a gain on the amplitude of the synaptic kernels. In effect, this parameterization controls the populations sensitivity to afferent inputs (Kiebel et al., 2007). For simplicity, we only considered fluctuations in extrinsic connectivity.

Following the inversion of the group PEB, a greedy search was performed to prune any parameters that do not contribute to the log evidence (i.e., parameters that increase complexity without increasing accuracy). The search algorithm used Bayesian model reduction (BMR) to remove redundant connection parameters from the full model; until there was no further improvement in model-evidence. The parameters of the best 256 models from this search procedure were then averaged, weighted by their model evidence (i.e. Bayesian Model Averaging, BMA). This procedure was performed for both networks. The results for the default mode network and the saliency network are shown in Fig. 4 and 5 respectively. From Fig. 4, it is clear that the DCT components for the DMN were either pruned away during the greedy search or that the posterior probability was below .90. More specifically, only for the first DCT component, three of the connectivity parameters survived Bayesian model reduction. Interestingly, only the constant term and the monotonic decay basis function showed a clear effect. For the monotonic decay, the forward connections showed a negative effect (resulting in monotonic increasing trajectory) and the lateral connections showed a positive decay effect. In order to demonstrate the time dependent effect on effective connectivity parameters more clearly, the (posterior predictive expectations of the) group level trajectories for each of the DCM parameters are shown in Fig.6A.

**Figure 4.**
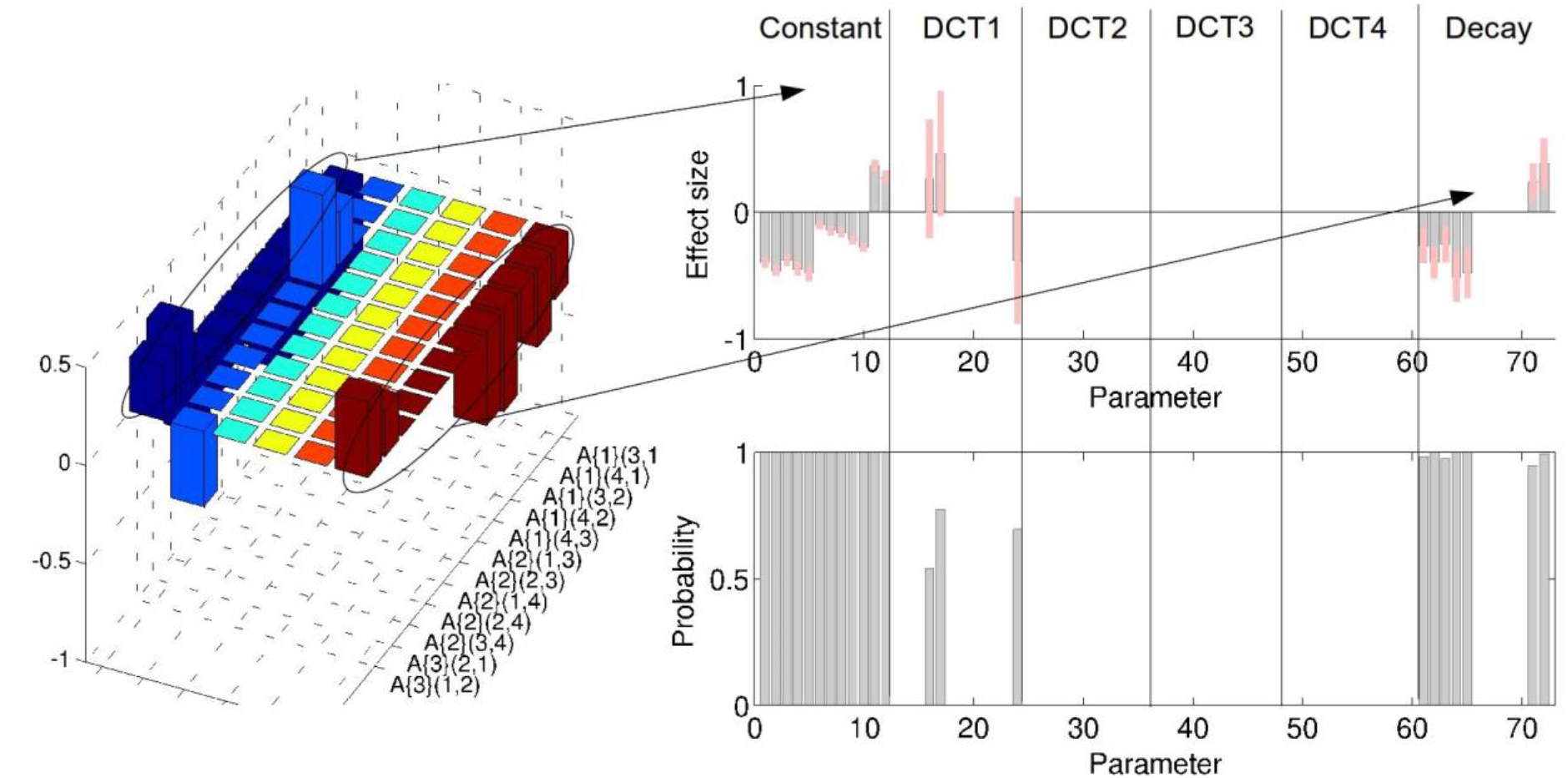
The results of the group PEB, following BMA for the default mode network. On the left side, a 3D bar graph reports all the parameters of the model. The key for the connectivity codes can be found in Table 1. On the top-right panel, the same parameters are shown together with a 90% Bayesian confidence interval. Bottom right panel shows the probability of each parameter in the group PEB. This posterior probability compares all models with and without the effect (i.e., parameter) in question.

**Figure 5.**
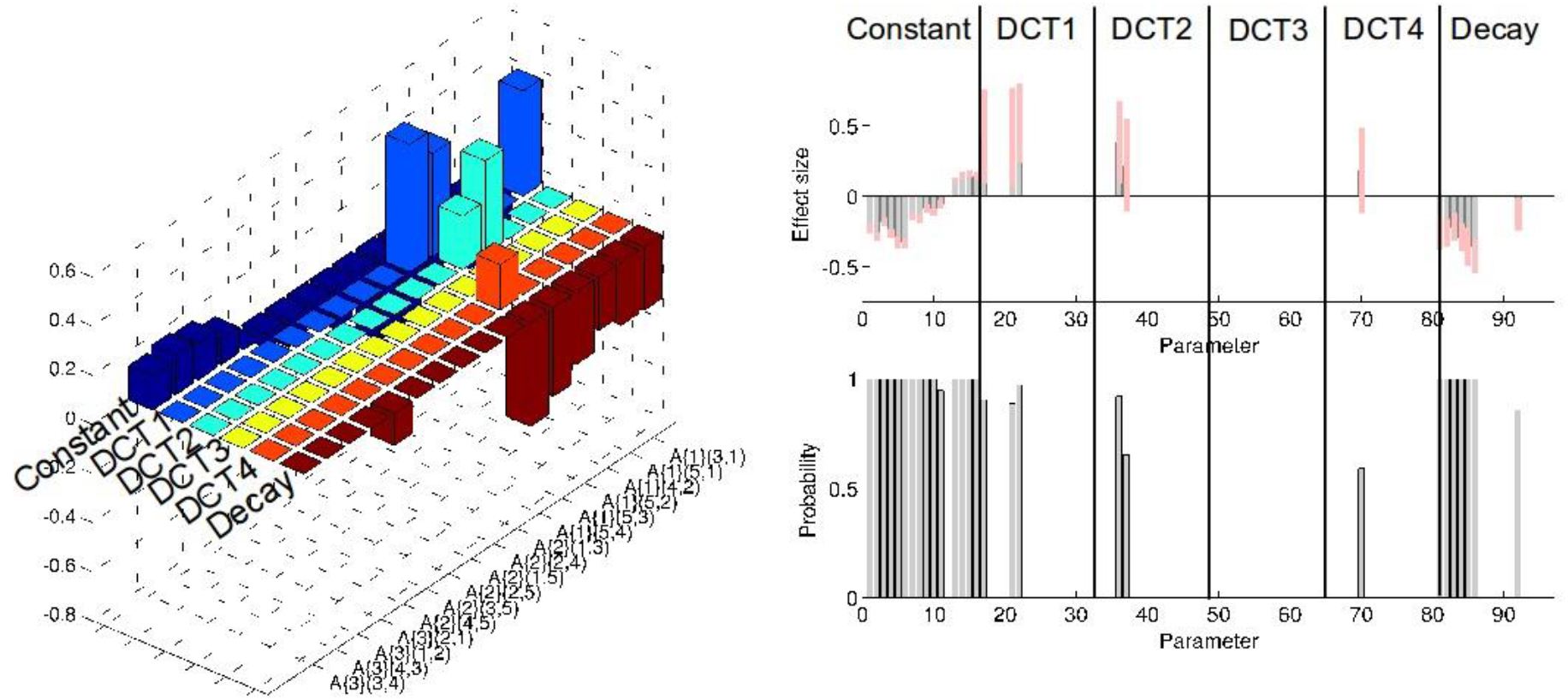
Same format as for Fig 4 but now for the saliency network.

**Figure 6.**
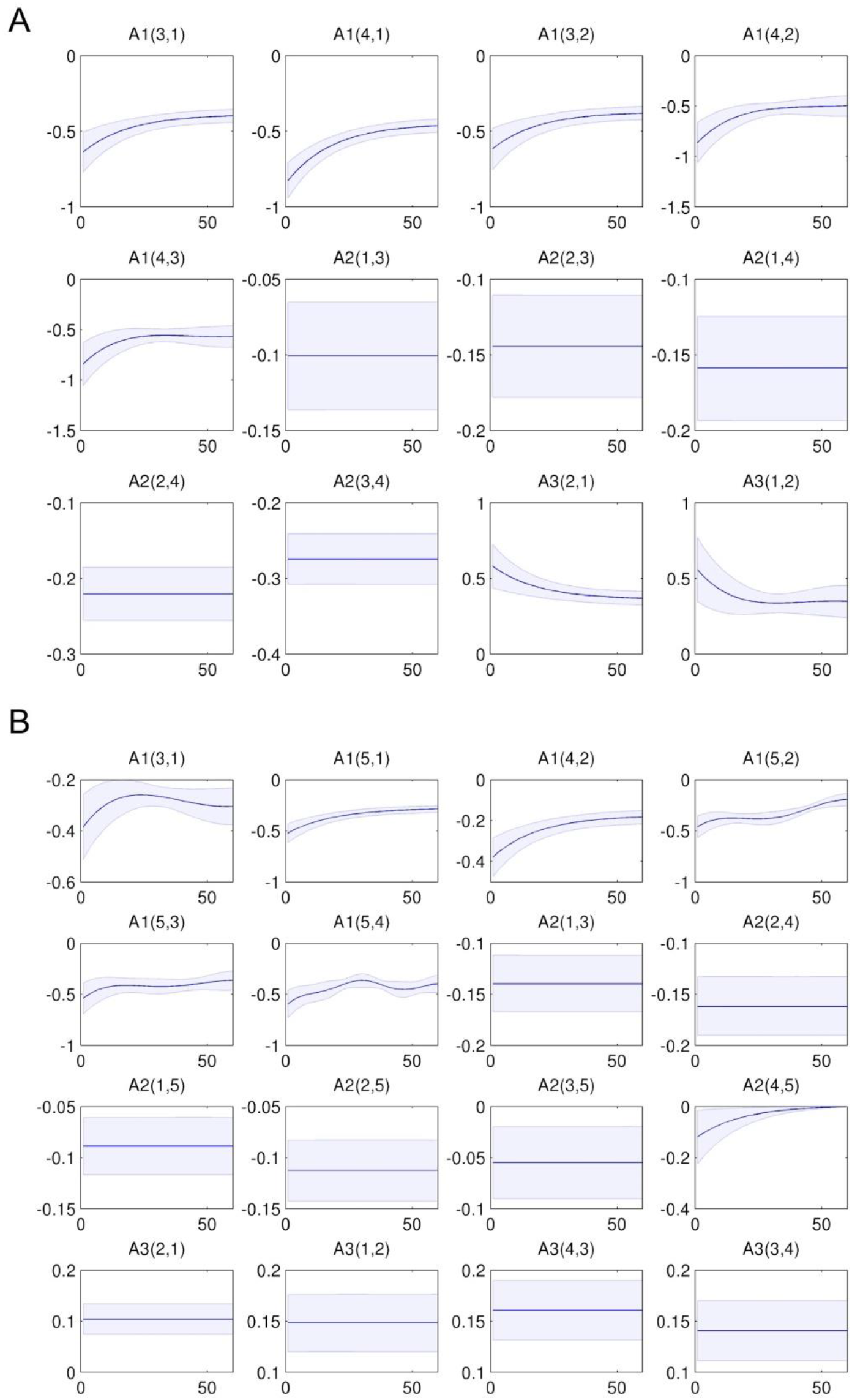
Predicted DCM (log scale) parameter trajectories for the default network (A) and for the saliency network (B). The blue shaded areas around the lines indicate a 90% confidence interval (i.e., Bayesian credible interval) of the predicted trajectories for each connection at the group level. A1, A2 and A3 refer to forward backward and lateral connections respectively.

In Fig. 5, the group PEB parameters after BMR and BMA are shown for the saliency network. For the decay component, a similar pattern was observed: all forward connections showed a negative monotonic decay effect. With respect to the DCT components, six parameters survived Bayesian model reduction, which in some cases showed an effect with relative high probability.

In Fig 6B, the predicted trajectory of each connection, at the group level, are shown for the saliency network. Here, the effects of the DCT components were more prominent compared to the default network. Nevertheless, the pattern is highly comparable with the default network; namely, a monotonic increase of the forward connections.

## Discussion

In this work, we build upon recent characterizations of dynamic brain connectivity at rest (e.g. Allen et al., 2014; Chang and Glover, 2010; Vidaurre et al., 2017; Wang et al., 2016). Here, we used the approach proposed recently by Park et al. (2017). Parametric empirical Bayes (PEB) was employed to model fluctuations in effective connectivity; where effective connectivity was inferred using dynamic causal modelling (DCM). One minute rest-EEG time-series were segmented into 60 non-overlapping windows. Then DCM for cross spectral densities was (iteratively) fitted to the data for each window. Afterwards the differences in DCM parameters between the windows were modelled with a second level linear model. The design of this second level comprised of a number of temporal basis functions plus a constant term. This way time dependent fluctuations in effective connectivity were modelled. Here, we used 4 DCT components and one exponential decay component – whose linear combinations can cover a wide range of plausible trajectories. This hierarchical model was estimated with PEB. The main advantage of PEB is that deep hierarchical models can be inverted recursively bypassing posterior densities to subsequent levels (i.e., PEB of PEB). This allowed us to ask which components of the parameter trajectories are conserved over subjects. Another advantage of PEB is that reduced second (or third) level models can be derived efficiently from a full model, without having to re-estimate the reduced lower levels. Note that, in this (EEG) study, changes in effective connectivity were investigated at much faster time scales compared to fMRI.

Bayesian model comparison of the second level (between-window, within-subject) models showed that for the default mode network, the model containing only the exponential decay basis function was the best out of 32 reduced models. For the saliency network, the model containing all basis functions appears to be the winning model. Thus, we provide evidence that effective connectivity parameters change systematically over time in both networks, even over the course of one minute using EEG. In other words, the observed differences between windows cannot be explained by non-systematic random fluctuations around a constant; when comparing the log evidence for second level models (which embodies a trade-off between model accuracy and model complexity).

Comparing the results for the default mode network with the results from Park et al. (2017) that used resting state fMRI, we see that in their analysis, all DCT components (for one or more DCM connectivity parameters) appear to be necessary to explain between window differences. Note that Park et al did not include an exponential decay function in their analysis. In the present dataset, subjects were asked to keep their eyes open for one minute, which was then followed by short block in which they were instructed to keep their eyes closed – and other movement task blocks). Therefore, subjects in our study might still be in a sort of adaptation phase during acquisition of the data. Park et al. (2017), on the other hand, used data from the Human connectome project, in which participants are asked to relax inside the MRI scanner for approximately 15 minutes. This difference in context might explain why our findings differ from Park et al. (2017).

Considering the group level parameters in Fig. 4 and 5 – and the group level estimates of the connectivity trajectory of both networks (Fig 6.) – the similarities are apparent. In both networks, we see that most log-scaling parameters are negative for both the forward and backward connections. In addition, the DCT components do not seem to have a substantial effect on connectivity fluctuations that are preserved over subjects. On the other hand, a reversed exponential decay for the forward connection is apparent in both networks. Looking at the group level trajectory, both networks give qualitatively similar temporal effects with respect to the forward and backwards connections. In the default mode network we see a monotonic decay of the lateral connection while no temporal effect on the lateral connection in the saliency network can be observed.

As noted above, negative log-scaling parameters mean that the fixed values corresponding to excitatory connections in the neural model are downscaled by the scaling parameter. In the neural mass model we used here, forward connections have a strong driving effect, while backward connections have more inhibitory and modulatory influence on the nodes they target. This difference in forward and backwards connections implies a functional cortical hierarchy. In light of our results, connectivity from low to higher order brain regions becomes more pronounced. This suggests that more information is conveyed through the network with time in ascending or bottom-up fashion during this first minute of rest eyes-open period.

Given the similarities of the group level trajectories between the networks, these results appear to be at odds with the results of the Bayesian model comparison of second level models. More specifically, the absence of many significant DCT components at the group level for the saliency network seems to be at odds with the BMS suggesting the contribution of all basis functions at the within-subject level. This apparent discrepancy can be resolved by considering that the PEB treats the DCM parameters (or second level parameters for the PEB of PEB) as random effects, with mean equal to (***X_j_*** ⊗ ***X_j_***)***β_j_***. Therefore, although at the individual subject level, the DCT components show significant effects; these are not conserved over subjects. In other words, they have a group mean of zero. This also means that the trajectories are unique for each individual. Only the decay component embedded in the parameter trajectories is conserved over subjects. For the default mode network however, the DCT effects at the individual subject level are not systematically different from zero. A possible explanation might be that subject specific factors – such as age, gender, and fatigue – had a larger effect on the saliency trajectories compared to the default mode network. For the default mode, the effect might be mainly driven by the context, which is common to all subjects. In sum, we showed that connectivity trajectories can contain components that are subject-specific but systematic (e.g. DCT components in the saliency network) and components which are conserved over subjects (e.g. monotonic increase in both networks).

Here, we heuristically chose a window size of 1 second. However, window size can have a significant effect on the resulting connectivity dynamics. Shakil et al. (2016) for example showed that the window size was one of the most important factors in the variability of FC applied to rs-fMRI. A small window size could, for example, lead to poor estimation of FC. In the context of fluctuations in effective connectivity, one heuristic one can adopt is to choose the window length at which the parameters can accurately be obtained. This requires simulating data in which the ground truth is known, in order to evaluate the minimum window length required in a number of plausible situations (size of network, number of channels etc.). In the context of DCM for rs-fMRI, this approach has been undertaken in (Razi et al., 2015). For EEG however, the situation remains unclear and should be further investigated.

Given the non-invasive nature, relative low cost and portability of EEG, tracking time-variability of connectivity can be readily applied in patient studies. Deriving connectivity fluctuations from EEG might be a promising avenue for future research. Differences in parameter trajectories could be used as a neural signature of pathophysiology or psychopathologies. Also, treatment effects could be related to connectivity fluctuations. Another possible application of the approach is to model factors such as fatigue, motivation, and learning over the course of an experimental procedure. This would require dividing the experiment, *post-hoc,* in a number of blocks in which trials are averaged (i.e. creating ERP’s). Although this would reduce the signal to noise ratio of the ERP’s within blocks, modelling such factors might outweigh the cost. For example, it could be that the mismatch negativity, a brain response to a violation of a rule, is reduced towards the end of an experiment. Using hierarchical modelling one could then disambiguate between e.g. low level reduction in sensitization and reduced top-down influences (see Garrido et al., 2009 for a similar approach). To conclude, our study showed that effective connectivity showed both common (saliency network and default mode network) and subject-specific (saliency network) trajectories over the course of 1 minute rest EEG.

## Acknowledgements

This research was supported by the Fund for Scientific Research-Flanders (FWO-V, grant No. FWO14/ASP/255 awarded to FVDS). The computational resources (Stevin Supercomputer Infrastructure) and services used in this work were provided by the VSC (Flemish Supercomputer Center), funded by Ghent University, FWO and the Flemish Government – department EWI. HA is funded by the Special Research Fund of Ghent University (awarded to HA; https://www.ugent.be/), KJF is funded by a Wellcome Trust Principal Research Fellowship (Ref: 088130/Z/09/Z).

